# Genome sequencing and comparative analysis of *Wolbachia* strain *w*AlbA reveals *Wolbachia*-associated plasmids are common

**DOI:** 10.1101/2022.07.01.498274

**Authors:** Julien Martinez, Thomas H. Ant, Shivan M. Murdochy, Lily Tong, Ana da Silva Filipe, Steven P. Sinkins

**Author notes:** **Corresponding authors:** Julien Martinez,; Prof. Steven P. Sinkins,.

## Abstract

*Wolbachia* are widespread maternally-transmitted bacteria of arthropods that often spread by manipulating their host’s reproduction through cytoplasmic incompatibility (CI). Their invasive potential is currently being harnessed in field trials aiming to control mosquito-borne diseases. *Wolbachia* genomes commonly harbour prophage regions encoding the *cif* genes which confer their ability to induce CI. Recently, a plasmid-like element was discovered in *w*Pip, a *Wolbachia* strain infecting *Culex* mosquitoes; however, it is unclear how common such extra-chromosomal elements are in *Wolbachia*. Here we sequenced the complete genome of *w*AlbA, a strain of the symbiont found in *Aedes albopictus*. We show that *w*AlbA is associated with two new plasmids and identified additional *Wolbachia* plasmids and related chromosomal islands in over 20% of publicly available *Wolbachia* genome datasets. These plasmids encode a variety of accessory genes, including several phage-like DNA packaging genes as well as genes potentially contributing to host-symbiont interactions. In particular, we recovered divergent homologues of the *cif* genes in both *Wolbachia*- and *Rickettsia*-associated plasmids. Our results indicate that plasmids are common in *Wolbachia* and raise fundamental questions around their role in symbiosis. In addition, our comparative analysis provides useful information for the future development of genetic tools to manipulate and study *Wolbachia* symbionts.

## Introduction

*Wolbachia* is the most abundant heritable bacterium in arthropods, present in around half of the species worldwide (Weinert et al. 2015), as well as being an obligate symbiont of filarial nematodes (Taylor et al. 2005). It is primarily inherited through the female germline and has evolved various ways to spread through host populations by manipulating arthropod reproduction. Reproductive alterations include several forms of sex-ratio distortions and cytoplasmic incompatibility (CI), a type of selective sterility providing a reproductive advantage to female hosts carrying the symbiont (Kaur et al. 2021). Many *Wolbachia* strains are also capable of inhibiting viral replication (Teixeira et al. 2008; Walker et al. 2011; Martinez et al. 2014; Ant et al. 2018) and this phenotype, combined with the self-spreading mechanism of CI, allowed the development of novel strategies for controlling mosquito-borne diseases (Nazni et al. 2019; Gesto et al. 2021; Utarini et al. 2021).

*Wolbachia* is found exclusively within the host cell environment, and this has hampered the use of genetic tools to manipulate and study its genome at the mechanistic level. Nevertheless, genome research has led to considerable progress in understanding *Wolbachia* biology (Wu et al. 2004; Chrostek et al. 2013; Ellegaard et al. 2013; Gerth et al. 2016). *Wolbachia* is commonly associated with prophage WO, a temperate bacteriophage integrated as a prophage region into the symbiont’s chromosome. Prophage WO has an important role in *Wolbachia*’s evolution by allowing the transfer of genetic material between symbiont genomes (Bordenstein and Bordenstein 2022). Prophage WO also carry an accessory region called the Eukaryotic Association Module (EAM) which encodes a variety of genes that are eukaryotic-like in length, origin and/or predicted function such as ankyrin domain-containing proteins. Moreover, the EAM often carries the two syntenic genes *cifA* and *cifB* responsible for the CI phenotype (Bordenstein and Bordenstein 2016; Beckmann et al. 2017; Lepage et al. 2017). While the mobility of phage WO has contributed to horizontal gene transfer over long evolutionary periods, evidence of active phage replication and production of virus particles remain limited to a few examples (Masui et al. 2001; Fujii et al. 2004; Bordenstein and Bordenstein 2016; Kupritz et al. 2021). In addition, many prophage regions display signs of degradation through pseudogenization and chromosomal rearrangements (Bordenstein and Bordenstein 2022).

Recently, a new kind of mobile genetic element was discovered in *w*Pip, a strain of *Wolbachia* infecting *Culex pipiens* mosquitoes (Reveillaud et al. 2019). The 9,228 bp extrachromosomal and circular element bears the hallmarks of a candidate plasmid and was therefore named pWCP for “plasmid of *Wolbachia* endosymbiont in *C. pipiens*”. Since this discovery, no other *Wolbachia*-associated plasmids has been described and it is unclear how common these elements are and whether they play an important role in *Wolbachia*-host interactions.

There is abundant literature highlighting the contribution of plasmids in the horizontal transfer of adaptive traits such as antibiotic resistance and virulence in free-living bacteria (Bennett 2008; Norman et al. 2009; Rodríguez-Rubio et al. 2020). Plasmids have also been described in arthropod symbionts and they often carry genes potentially involved in symbiosis. For instance, ankyrin repeat-containing and toxin-like genes are commonly found in the plasmids of *Cardinium* (Penz et al. 2012; Santos-Garcia et al. 2014), *Spiroplasma* (Masson et al. 2018; Pollmann et al. 2022) and *Rickettsia* symbionts (Gillespie et al. 2014). In *Spiroplasma* strain MSRO infecting *Drosophila melanogaster*, a male-killing phenotype is mediated by the *spaid* toxin encoded on plasmid pSMSRO (Harumoto and Lemaitre 2018). Homologues of the *Wolbachia cif* genes have also been identified on plasmid pLbaR in *Rickettsia felis*, however, their role in altering host reproduction is not established (Gillespie et al. 2018; Martinez et al. 2021).

The invasive ‘Asian tiger’ mosquito *Aedes albopictus*, a vector of arboviruses such as dengue, chikungunya and Zika, harbours two co-infecting strains of *Wolbachia* called *w*AlbA and *w*AlbB. *w*AlbA has a lower overall density and in females is largely restricted to the ovaries, while *w*AlbB has a somewhat wider tissue distribution (Dutton and Sinkins 2004; Ant and Sinkins 2018). Following transfer into the naturally *Wolbachia*-free *Ae. aegypti*, much wider tissue distribution occurs for both strains in this novel host, but their relative density was reversed; however, despite its lower density *w*AlbB was a much more efficient inhibitor of arbovirus transmission in *Ae. aegypti* than *w*AlbA (Ant et al. 2018). Subsequently *w*AlbB has been deployed in Malaysia for dengue control (Nazni et al. 2019). Several genomes of *w*AlbB have been sequenced (Sinha et al. 2019; Scholz et al. 2020; Ross et al. 2021), but due to the technical difficulties associated with its lower density and co-presence of *w*AlbB, no *w*AlbA genome has been reported to date.

Here we report the complete genome of *Wolbachia* strain *w*AlbA and two new associated plasmids, together with a number of similar plasmids and chromosomal islands in existing *Wolbachia* genome assemblies.

## Results

### *w*AlbA harbours prophage regions and multiple pairs of *cif* genes

In order to sequence the *w*AlbA genome, we generated a singly-infected *Ae. albopictus* line by eliminating the co-infecting strain *w*AlbB (see methods). We found that *w*AlbA is most abundant in the ovaries and its tissue distribution is not affected by the presence of *w*AlbB (Figure S1A). In addition, *w*AlbA does not affect the fecundity of female mosquitoes and is able to induce CI (Figure S1B-C). However, *w*AlbA is unable to rescue CI in crosses with doubly-infected males, suggesting that *w*AlbA and *w*AlbB carry incompatible pairs of *cif* genes (Figure S1C).

We assembled a 1,190,930 bp circular *Wolbachia* genome from *w*AlbA-infected ovaries (Figure 1A) which is similar in size and gene number to other complete *Wolbachia* genomes (Table S1). *w*AlbA belongs to the arthropod-specific *Wolbachia* supergroup A (Figure 2) and harbours a large WO prophage region (47,475 bp, Figure 1A) comprising a complete set of structural and non-structural gene modules thought to be essential for the production of phage particles (Table S2). However, this region displays a sequencing depth similar to the rest of the chromosome, suggesting no active replication of WO phage (Figure 2A). We also found smaller WO phage islands containing core phage genes, often with signs of pseudogenization, and accessory genes commonly found in the phage eukaryotic association module (Figure 1A, Table S2). This includes three nearly identical pairs of the cytoplasmic incompatibility genes *cifA* and *cifB*, all harbouring a *cifB* deubiquitinase domain that is characteristic of Type I *cif* homologues (Martinez et al. 2021).

**Figure 1.**
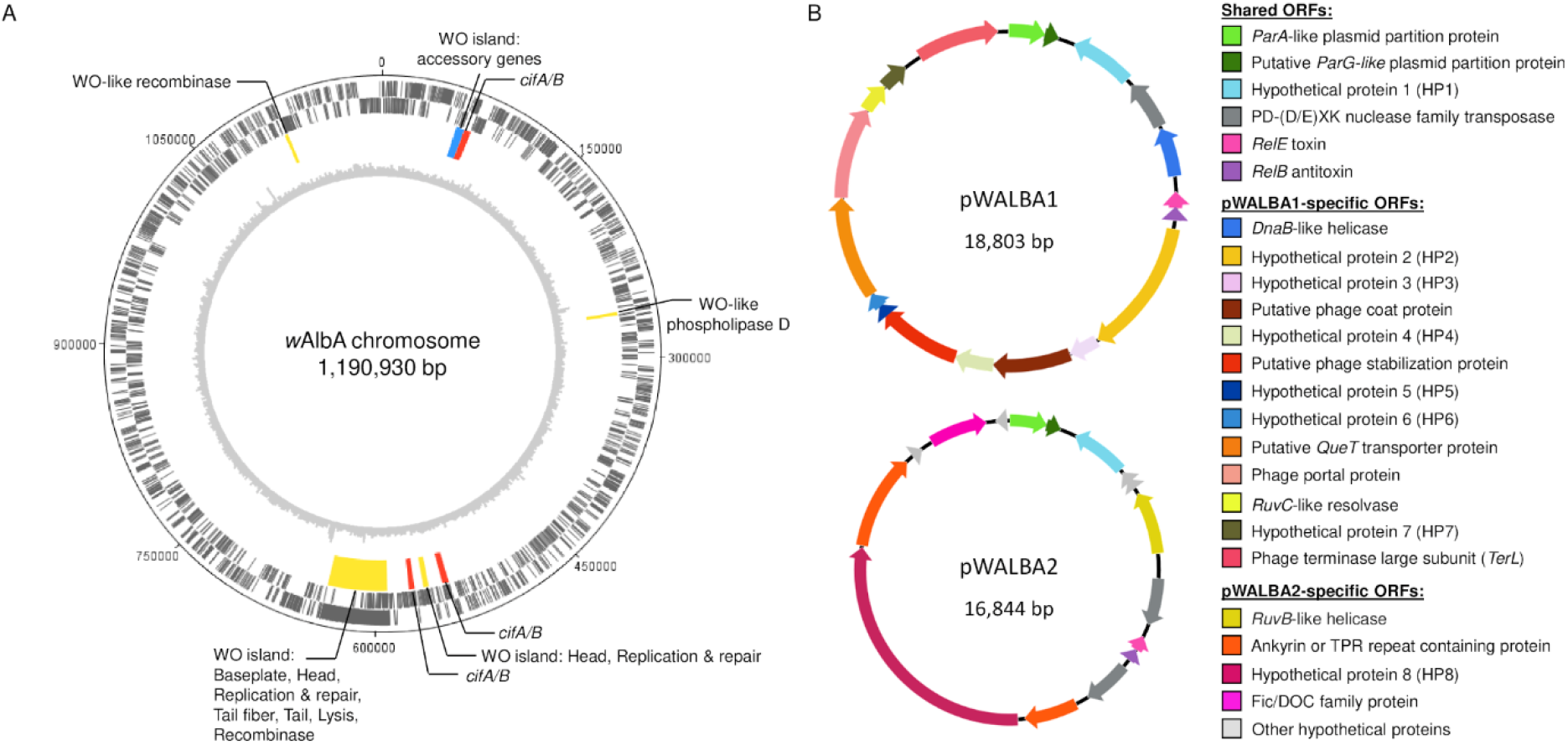
Genome map of the *w*AlbA chromosome and plasmids. (A) *w*AlbA chromosome. Two outer circles: protein-coding genes on forward and reverse strands (black). Third circle: WO prophage regions (yellow: phage core genes, red & blue: phage eukaryotic modules). Inner grey circle: Illumina sequencing depth per 2,000 bp window from a non-WGA mosquito sample (see methods). (B) *w*AlbA plasmids and predicted coding sequences.

**Figure 2.**
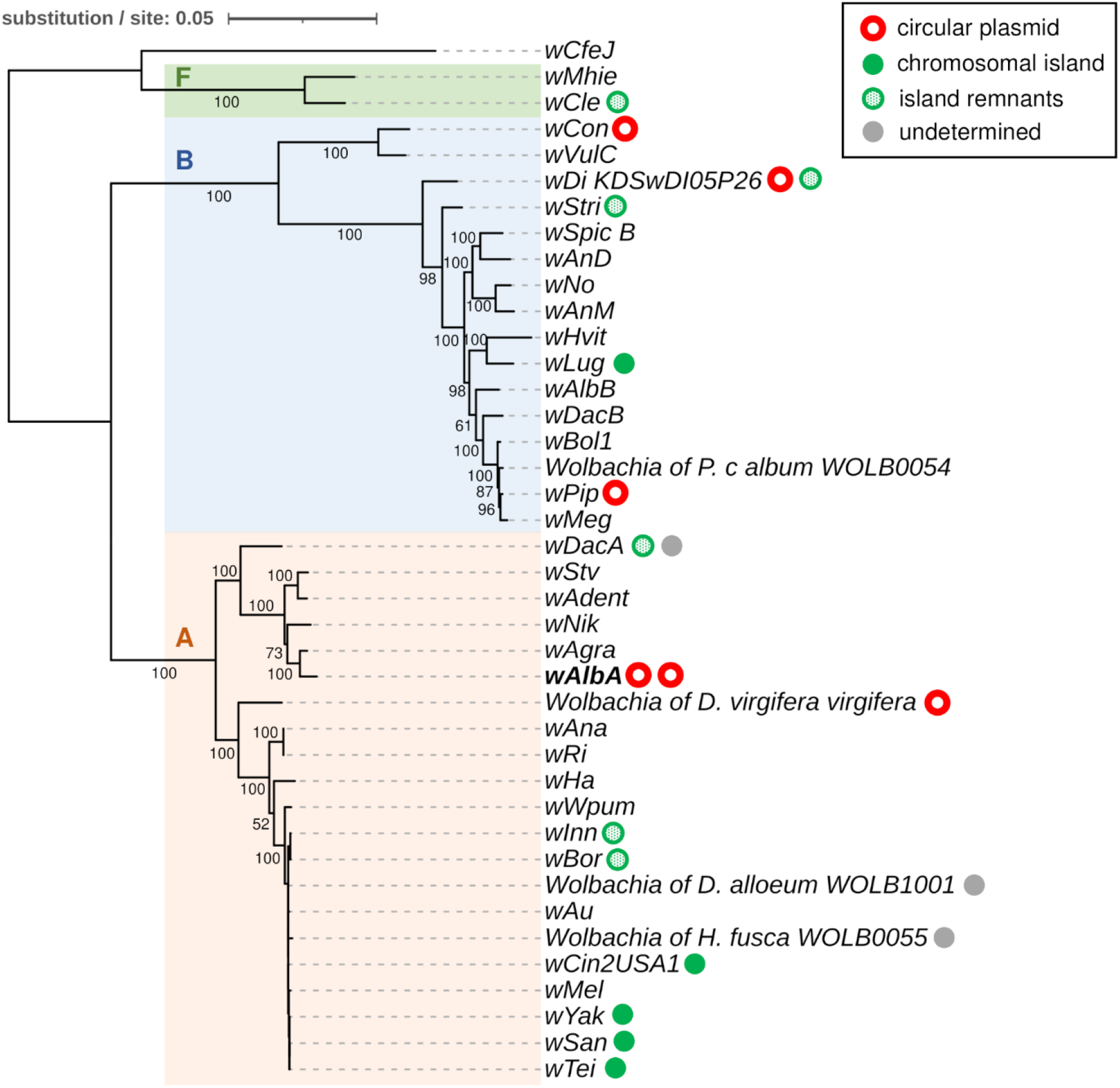
*Wolbachia* strain phylogeny and distribution of plasmid-like elements. Maximum Likelihood phylogeny based on the concatenated nucleotide alignment of 36 *Wolbachia* core genes. Branch support was assessed with 1,000 bootstrap replicates. Capital letters indicate *Wolbachia* supergroups. Circles indicate strains in which plasmid-like elements were identified.

### *w*AlbA is associated with two plasmid-like elements

In addition to the *w*AlbA genome, we assembled two circular extrachromosomal elements, 18,803 and 16,844 bp in size, that show strong similarities to pWCP, the *Wolbachia*-associated plasmid found in *C. pipiens* (Figure 1B). Based on the comparative analysis below, we named these plasmids pWALBA1 and pWALBA2 (for plasmids of *Wolbachia* endosymbiont *w*AlbA 1 and 2). Using specific PCR primers, we confirmed the association of the two plasmids with *w*AlbA, whereas no amplification was observed in the *Ae. albopictus* Aa23 cell line infected with *w*AlbB only (Figure S2). pWALBA1 showed a higher copy number per *Wolbachia* cell than pWALBA2, although pWALBA2 copies tended to increase in older female mosquitoes (Figure S3A). Importantly, the relative amounts of the two plasmids per mosquito were strongly correlated with *w*AlbA density, further supporting their association with the symbiont strain (Figure S3B).

Similar to pWCP, both *w*AlbA plasmids encode a *ParA*-like plasmid partitioning gene (HHpred probability > 99%), a hypothetical protein with strong similarity to a *ParG*-like plasmid partition gene and other DNA-binding proteins found on extra-chromosomal elements (e.g. phage transcriptional *Arc* repressor, HHpred p > 99%), a hypothetical protein (HP1), a *RelB/E* toxin-antitoxin addiction module (HHpred p > 98%) as well as one or two transposases (Figure 1B, Table S3). pWALBA1 shares additional genes with pWCP, namely a *DnaB-like* helicase (HHpred p = 99.96%) and a *TerL-like* phage terminase large subunit (HHpred p = 100%), but contrary to pWCP, these two genes do not show sign of pseudogenization. pWALBA1 encodes other phage-like proteins, namely a portal protein (HHpred p = 100%), a putative phage stabilization protein (HHpred p = 99.44%) and a hypothetical protein with weak homologies to a phage coat protein (HHpred p = 64.73%, Table S3). Other pWALBA1 genes include a putative *QueT*-like queuosine transporter (HHpred p = 95.17%), a *RuvC-like* resolvase (HHpred p = 98.17%) and several hypothetical proteins absent in pWCP. We did not find phage-like genes in pWALBA2 but instead genes encoding a Fic/DOC family protein (HHpred p = 100%), a *RuvB*-like helicase (HHpred p = 99.02%) and other proteins with no predicted functions, some of which carry ankyrin or tetratricopeptide (TPR) repeat domains (Figure 2B, Table S3).

### Plasmids and related chromosomal islands are widespread in *Wolbachia*

In order to investigate how common plasmid-like elements are in *Wolbachia*, we conducted TBLASTN searches of pWALBA1 and pWALBA2 coding sequences in publicly available *Wolbachia* genomes. We found plasmid-like regions with similar gene organization to *w*AlbA plasmids in 47 out of 189 *Wolbachia* assemblies (~20% of the strains, several assemblies for some strains) and none were found in strains from nematodes (Table S4). Their distribution is patchy across the *Wolbachia* phylogeny, suggesting that plasmids are often horizontally-transferred between symbiont strains and tend to be lost over long evolutionary timeframes (Figure 2). However, it is possible that plasmid sequences are missing from some *Wolbachia* assemblies. Indeed, filtering out contigs from final assemblies based on a lack of homology with known *Wolbachia* sequences or due to large differences in sequencing depth is common practice. For instance, not all draft assemblies of the *Wolbachia* infecting *Diabrotica virigifera virgifera* contained plasmid-like sequences (Table S4). Moreover, we found a plasmid-like contig in several isolates of strain *w*Di (Pascar and Chandler 2018) that was absent in other published *w*Di assemblies (Surendra et al. 2022). However, by re-assembling the raw reads from the latter study, we retrieved the corresponding circular plasmid genome that had not been reported (isolate *w*Di_KPSwDI05P26). We circularized additional plasmid-like elements from the *w*Con strain (SRA accession: SRR7213553), from a *Wolbachia-infected* sample of the *Diabrotica virgifera virgifera* beetle (SRA accession: SRR1106544) and from a sample labelled *Insecta*_WOLB1166 (mixture of *Wolbachia*-infected insect species, SRA accession: SRR748268). All circular plasmids show higher sequencing depth relative to their respective *Wolbachia* genome, ranging from ~4 to 10x, suggesting that they are maintained as multiple copies within bacterial cells (Figure 3). In some cases, copy number could not be estimated due to raw sequencing data being unavailable (*w*DacA) or to the presence of several *Wolbachia* strains in the same sample as indicated by multiple peaks in the sequencing depth distribution (*Insecta*_WOLB1166, *Dactylopius coccus* WOLB1009 and *Megaselia abdita* WOLB1013).

**Figure 3.**
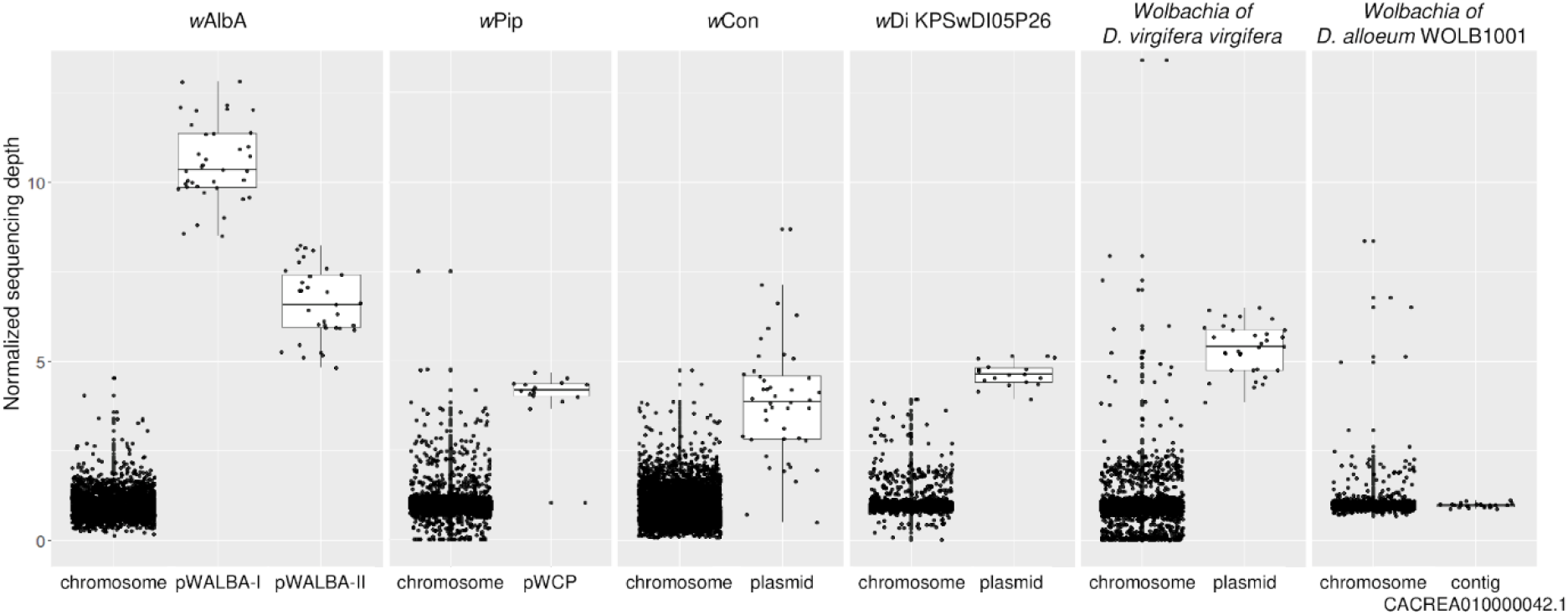
Relative sequencing depth of *Wolbachia*-associated plasmid-like elements. Each dot indicates the mean sequencing depth in 500 bp windows divided by the mean chromosome depth.

Some linear contigs could not be circularized but show a similar sequencing depth relative to other *Wolbachia* contigs and could be single-copy plasmids or be part of the bacterial chromosome (e.g. in *Wolbachia* of *Diachasma alloeum* WOLB1001, Figure 3). In support of the latter hypothesis, we found several genomic regions similar to pWALBA1 integrated on the *Wolbachia* chromosome (Figure 4). These chromosomal islands include a twelve gene region previously named “Dozen island” present in *Wolbachia* strains from *Drosophila teissieri, santomea* and *yakuba* that encompasses the variable region of pWALBA1-like plasmid genomes (Baião et al. 2021). In most cases, these islands are flanked by transposable elements and show varying degrees of chromosomal rearrangements and degradation with genes carrying premature stop codons or being truncated by transposase sequences (Figure 4). An interesting observation is that plasmid-like islands in strains *w*Oegib-WalA and *w*Oegib-WalA found in *O. gibbosus* spiders are located next to WO prophage regions (Figure 4).

**Figure 4.**
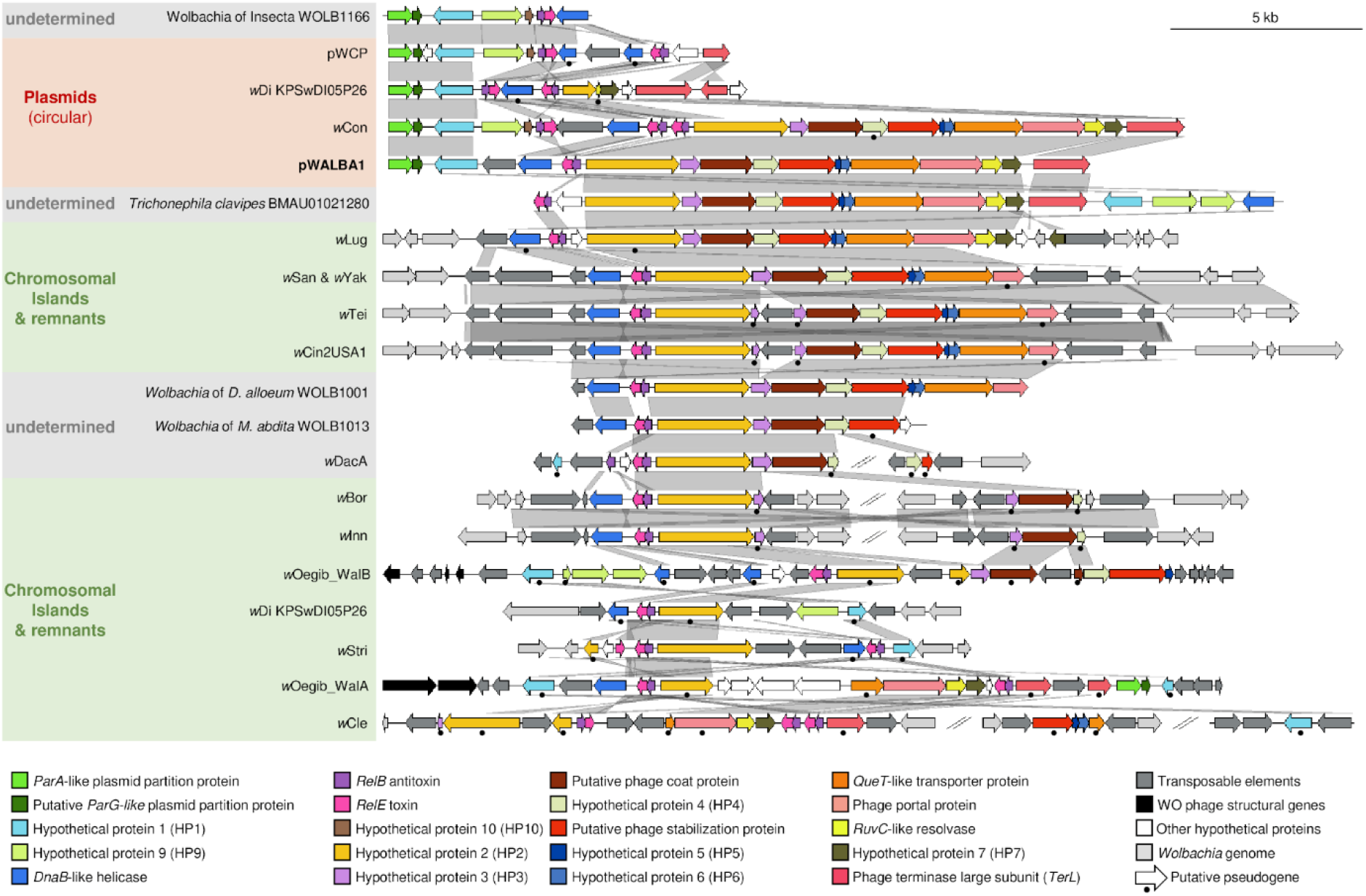
Gene organisation and comparison of pWALBA1-like sequences. Similarity is indicated by gene colours (TBLASTN) and by the grey areas between sequences (BLASTN) where darker grey means more similar.

Other circular plasmid sequences, including pWALBA2, were found to share the same set of core genes with the plasmids described above (*ParA-like, ParG*-like, HP1, *RelB/E*) while encoding different genes in their accessory region (Figure 5). These include a Fic/DOC family protein and other hypothetical proteins carrying ankyrin, PD-(D/E)XK nuclease and RDD domains. In addition, the plasmid found in the *Insecta_WOLB1166* sample encodes a *cifA/B* gene pair and the one from *D. virgifera virgifera* a *cifB* homologue lacking the 5’ end of the gene (Figure 5). In both cases, *cifB* harbours an ankyrin-repeat region as well as a C-terminal latrotoxin domain which was only found previously in divergent Type V *cifB* homologues (Figure 5) (Martinez et al. 2021).

**Figure 5.**
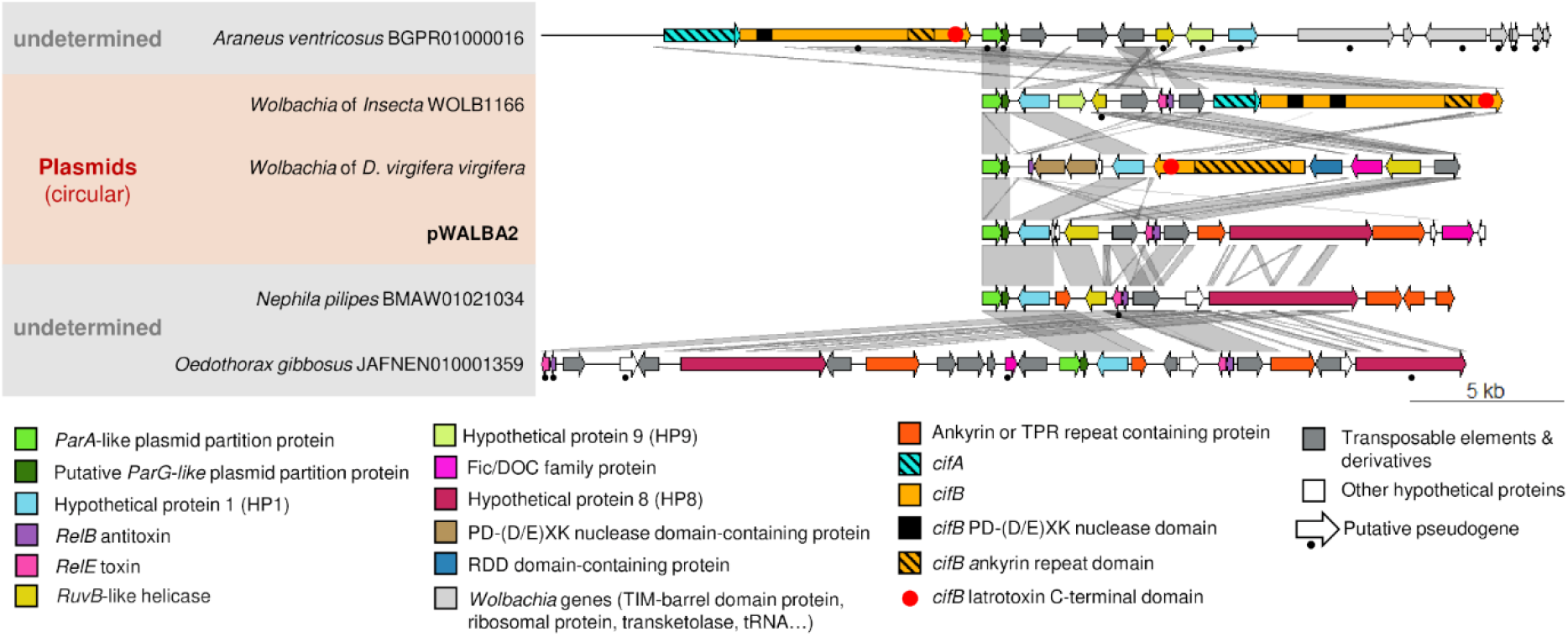
Gene organisation and comparison of other plasmid-like sequences. Similarity is indicated by gene colours (TBLASTN) and by the grey areas between sequences (BLASTN) where darker grey means more similar.

Plasmid-like sequences were also identified in draft genome assemblies of several species of spider (*Trichonephila clavata, T. clavipes, Araneus ventricosus, Nephila pilipes, Oedothorax gibbosus*) and one in the parasitic wasp *Cotesia glomerata* (Table S4, Figure 4 and 5). Some of these sequences are made of a short contig and could be *Wolbachia*-associated plasmids or chromosomal islands that were not filtered out from the host genome assembly. Indeed, *O. gibbosus* is known to host two *Wolbachia* strains (Halter et al. 2022) and we often found large contigs in these assemblies with strong homologies to *Wolbachia* genomes across their entire length (data not shown) suggesting the presence of the symbiont in these hosts. Nevertheless, two large spider contigs (*A. ventricosus* contig BGPR01000016.1, *T. clavipes* contig BMAU01021280.1) only showed homologies to *Wolbachia* in the region overlapping the plasmid-like sequences and could be cases of *Wolbachia*-to-spider horizontal transfers (Figure S4). However, we observed large variation in sequencing depth in the regions flanking the two putative plasmid inserts and low numbers of long Nanopore reads supporting these insertions (Figure S4). Therefore, we conclude that these are most likely chimeric contigs generated during the genome assembly of a *Wolbachia*-infected sample.

### Pervasive gene flow between *Wolbachia* and plasmid genomes

Phylogenetic analysis of the genes found in *Wolbachia* plasmids and chromosomal islands revealed the dynamic nature of lateral gene transfer and recombination between plasmids and the bacterial genome. Genes located on chromosomal islands do not form a monophyletic group which indicates that plasmid sequences have integrated into *Wolbachia* genomes more than once (Figure S5-S8). The phylogenetic relationships of genes in the *Dozen* island found in the *w*Tei-*w*Yak-*w*San clade and other closely-related *Wolbachia* strains suggest, however, a single origin of this island that has most likely codiverged with the bacterial chromosome following its integration. Genes of the *Dozen* island are on average more similar to homologues found in pWALBA1, while genes located on other islands such as in *w*Lug and *w*Stri tend to be more closely-related to the *w*Con plasmid (Figure S5-S7). The opposite pattern was found for the putative coat protein where the *w*Con homologue is more similar to its counterparts located within the *Dozen* island suggesting a possible recombination event. Another case of a probable recombination is observed between pWALBA1 and pWALBA2 since they carry divergent *ParA*-like plasmid partition genes while their genes encoding HP1 are closest relatives (Figure S5).

Several plasmid genes also have closely-related homologues on the bacterial chromosome located outside of plasmid-like islands, although we cannot rule out that they are island remnants. In the case of *ParA*-like homologues, chromosomal versions of the gene form a separate clade and, interestingly, they are all fused with the downstream *ParG-like* gene, as opposed to plasmid-encoded homologues (Figure S5). For other genes, there is evidence of multiple horizontal transfers between plasmids and *Wolbachia* genomes, including with WO prophage regions. RelB/E addiction modules are widespread in *Wolbachia* genomes (Fallon 2020) and their phylogenetic distribution does not suggest single origins for chromosomal, WO phage and plasmid homologues but rather multiple transfers between the different genomic entities (Figure S9). Similarly, the two PD-(D/E)XK nucleases located on the *Wolbachia* plasmid in *D. virgifera virgifera* are related to different lineages of nucleases recently described as being part of some WO phage head modules (Fallon 2022) (Figure S8). Finally, the plasmid Fic/DOC family proteins of pWALBA2 and *D. virgifera virgifera* group with two different clades of chromosomal Fic proteins commonly found in *Wolbachia* genomes, again suggesting multiple gene transfers between plasmids and *Wolbachia* genomes (Fallon 2020) (Figure S8).

### Are plasmids vectors of *cif* genes across the *Rickettslales?*

The *cif* genes are most of the time associated with WO prophage regions in *Wolbachia*. Previous studies have delimited multiple *cif* clades denominated as Type I to IV and a proposed Type V comprising homologues carrying additional domains on the C-terminal part of the *cifB* protein. *cif* homologues have also been reported in other *Rickettsiales*, some of them carried by plasmids (Gillespie et al. 2018; Davison et al. 2021; Martinez et al. 2021; Takano et al. 2021). We investigated the evolutionary origin of the *cif* genes located on *Wolbachia*-associated plasmid sequences and found that they group with Type V homologues (Figure 6), which is consistent with the length and predicted domains of their *cifB* protein (Figure 5). Unlike Type I-IV homologues, Type V *cif* genes are composed of several divergent clades with both *Wolbachia* and non-*Wolbachia* homologues and may not form a single monophyletic group (Figure 6). While the exact position of the tree root is currently unknown, we found evidence for several horizontal transfers of Type V *cif* genes between *Wolbachia* and other arthropod symbionts (Figure 6). Interestingly, the *cif* genes found in *Wolbachia*- and *Rickettsia*-associated plasmids do not cluster together suggesting independent acquisitions of *cif* genes by plasmids. In addition, we often found genes of the conjugation machinery in the regions flanking non-*Wolbachia cif* genes. In the chromosome of *Orientia tsutsugamushi* and the *Rickettsia* symbiont Oegib-Wal of the spider *O. gibbosus*, the *cif* genes are located next to an Integrative Conjugal Element (ICE), a type of mobile genetic element commonly found in *Rickettsia* genomes and plasmids (Akayama et al. 2008; Gillespie et al. 2014) (Figure 6; Figure S10). Other non-*Wolbachia cif* genes were located on short contigs but we often found genes of the conjugative machinery in the flanking regions suggesting that they are also associated with a chromosomal ICE or a conjugative plasmid (Figure 6).

**Figure 6.**
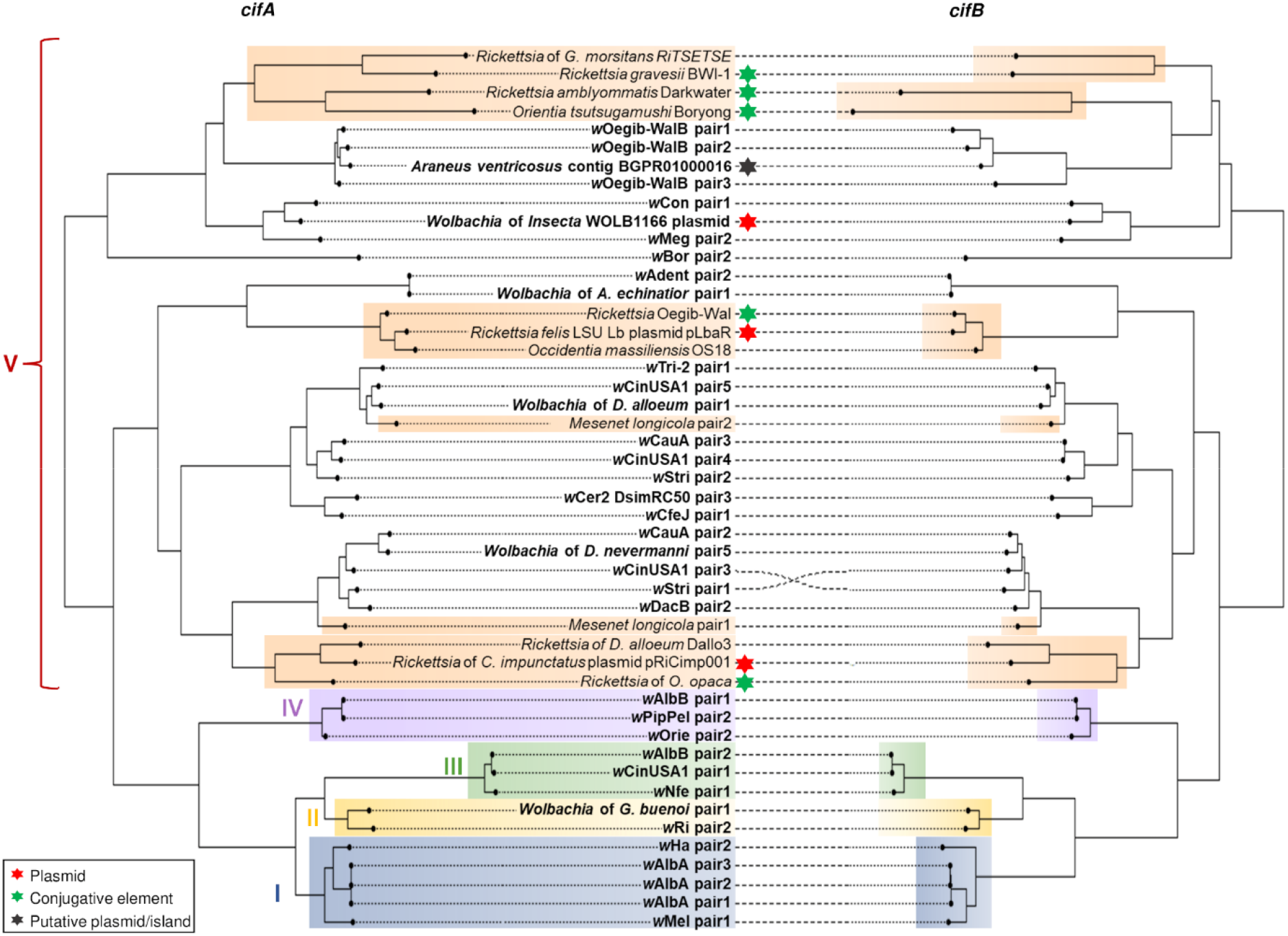
*Cif* gene phylogeny. Maximum Likelihood phylogeny based on the nucleotide alignment of *cifA* (left) and their cognate *cifB* genes (right). The tree is midpoint rooted. Previously defined *cif* Types I-IV are labelled and colour-shaded (blue/yellow/green/purple). Orange shading indicate Type V-like homologues found outside *Wolbachia* symbionts. *Wolbachia* homologue labels are in bold.

## Discussion

Until the recent discovery of the pWCP element (Reveillaud et al. 2019), it was assumed that plasmids were either absent or extremely rare in *Wolbachia*. Our study suggests that instead, these elements have been largely overlooked, likely as a consequence of them being discarded as contaminants or assumed to be part of the bacterial chromosome in *Wolbachia* sequencing projects. Our characterization of plasmids associated with *w*AlbA and other *Wolbachia* strains will facilitate the identification of related elements in future studies and should encourage the reanalysis of raw sequencing data available from public repositories.

In *w*AlbA, pWALBA1 and pWALBA2 were maintained as multiple copies per *Wolbachia* cell, however, pWALBA1 copy number was relatively stable while pWALBA2 copy number increased with the age of female mosquitoes. Sequencing depth analysis indicated that other *Wolbachia*-associated plasmids also tend to be maintained as multi-copy plasmids, similar to what was observed in (Reveillaud et al. 2019). We found large variation in size among the complete circular plasmid genomes, ranging from ~9,000 to 21,000 bp, which is primarily driven by differences in the presence/absence and pseudogenization of accessory genes. Nevertheless, all plasmids appear to share a set of core genes likely to be essential to their maintenance. The *ParA*-like and downstream *ParG*-like genes have strong structural similarities with proteins involved in plasmid partition systems that ensure plasmid inheritance during cell division (Baxter and Funnell 2014). *ParA* family proteins are ATPases that drive the segregation of newly replicated plasmids into daughter cells (Radnedge et al. 1998) while *ParG* is known to act as a centromere-binding protein modulating the action of its cognate ATPase (Wu et al. 2011). All plasmids also carry one or several *RelB/E* addiction modules which are Type II toxin-antitoxin (TA) systems that can promote plasmid maintenance through post-segregational killing of plasmid-free daughter cells (Gotfredsen and Gerdes 1998). *RelB/E* modules are also widespread in bacterial chromosomes, including in *Wolbachia* within and outside of prophage regions (Fallon 2020; Fraikin et al. 2020). The role of chromosomal TA systems is still being debated but they have been implicated in various functions such as stress-response (Christensen et al. 2001), anti-phage defences (Hazan and Engelberg-Kulka 2004) and may also prevent the loss of large genomic regions under fluctuating selection regimes (Szekeres et al. 2007). *RelB/E* modules may also accumulate as selfish DNA in bacterial chromosomes without providing fitness benefits due to their addictive nature (Fraikin et al. 2020).

A striking feature of some of the *Wolbachia* plasmids is the presence of several phage-like genes raising the possibility that they produce virus particles and could be transmitted horizontally via transduction. The two DNA packaging genes encoding the terminase large subunit and portal protein may allow plasmid genomes to be incorporated into viral capsids, while the presence of a phage stabilization and a putative coat protein indicate that plasmids might encode their own capsid structural proteins. We did not detect phage tail genes; however, such genes could be among the hypothetical proteins present on the plasmids or tail-less virus particles could be produced instead. Another hypothesis is that plasmids may highjack the tail proteins of WO phage, although this is perhaps unlikely since plasmid-encoded DNA packaging and structural phage-like proteins are not closely-related to WO homologues. It is however interesting to note that several plasmid genes have homologues in WO prophage regions (addiction modules, *cif* genes, PD-(D/E)XK nuclease) and that in two *Wolbachia* genomes, plasmid-like islands are located next to phage regions suggesting that plasmids may sometimes replicate and/or be packaged along with WO phage genomes. The existence of extrachromosomal phages, maintained as circular plasmids has been known for decades (Łobocka et al. 2004; Utter et al. 2014; Gilcrease and Casjens 2018) and a recent computational analysis revealed that such phage-plasmids are widespread in bacteria and encompass genetically diverse groups of mobile genetic elements (Pfeifer et al. 2021). Whether *Wolbachia*-associated plasmids produce phage particles remains to be investigated. While being genetically related, not all *Wolbachia* plasmids carry phage-like genes. This could mean that they rely solely on vertical transmission as we did not find that they encode their own conjugative machinery. There is evidence that some plasmids and phage-inducible chromosomal islands can undergo transduction by hijacking and even remodelling the capsid of coinfecting bacteriophages (Penadés and Christie 2015; Fillol-salom et al. 2019; Humphrey et al. 2021). Thus, it is possible that plasmids not carrying phage genes behave like phage satellites by being packaged into virions produced by WO prophage regions or by another coinfecting plasmid such as pWALBA1.

Over long evolutionary times, some *Wolbachia*-associated plasmids become integrated into the bacterial chromosome, a process that could be facilitated by the presence of many transposase sequences in plasmids and *Wolbachia* genomes. It also appears that both plasmids and chromosomal islands rapidly go extinct as indicated by their patchy distribution among *Wolbachia* strains and the abundance of pseudogenes. Their long-term persistence may therefore depend on a positive balance between new acquisitions through horizontal transfers of plasmids and losses through intense pseudogenization prior to or following an integration into the bacterial chromosome. The presence of RelB/E addiction modules in plasmids and islands likely contributes to lower the speed at which they become lost by preventing large deletions and plasmid losses during cell division. Selection may also act to maintain some of the plasmid/island genes if they confer benefits to *Wolbachia*. In particular, it is of interest that some *cif* genes are found on plasmids in both *Wolbachia* and *Rickettsia*. Although the function of plasmid-encoded *cif* proteins remains to be tested experimentally, they may provide the ability to induce CI and therefore contribute to the spread of the symbiont through host populations. There may even be an advantage for the symbiont in carrying *cif* genes on multi-copy plasmids rather than in the chromosome if an increase in the expression level of the *cif* proteins can modulate the CI phenotype and its rescue. Importantly, we showed that beyond phage WO, the *cif* genes can be associated with other mobile genetic elements like plasmids and integrative conjugal elements which likely contributed to their early evolution and spread across alphaproteobacterial symbionts.

In conclusion, our discovery of new plasmids associated with *w*AlbA and other *Wolbachia* strains provides a new framework for studying the role of these mobile genetic elements in *Wolbachia*. Importantly, our comparative analysis will guide future attempts aiming to develop a genetic tool kit for *Wolbachia* transformation. In particular, we expect that successful plasmid constructs will likely require some of the core genes we characterized such as those of the plasmid partition system and addiction modules.

## Material and methods

### Generation of a *w*AlbA-infected mosquito line

The *Ae. albopictus* wild type strain used in this study was collected from the Jalan Fletcher area of Kuala Lumpur (JF), Malaysia, in 2017. To generate a *w*AlbA-single infection, wild-type (*w*AlbA/*w*AlbB-carrying) larvae were reared under conditions of heat stress (diurnal rearing temperatures of 32-37°C). The resulting adult females were mated en-masse to males of a previously tetracycline-treated *Wolbachia*-negative JF line and were blood-fed and individualised for oviposition. An isofemale line carrying only *w*AlbA was isolated. Females from this line were subsequently outcrossed to *Wolbachia*-negative males from the JF line for five consecutive generations before a stable self-crossed colony was established.

*Wolbachia* density was measured in the dissected tissues of 5-day old adult females by quantitative PCR (qPCR) using *w*AlbA-specific primers (Table S5) normalised to the host homothorax (HTH) gene using a SYBR Green qPCR master mix (Bimake), with the following conditions: 95°C for 5 mins, followed by 40 cycles of 95°C for 15 secs and 60°C for 60 secs, and subsequent melt curve analysis.

Fecundity of females of the wild-type, *w*AlbA-only and *Wolbachia*-negative lines was measured by crossing females of each line to *Wolbachia*-negative males in pools of 10 males to 5 females, followed by blood-feeding and selection of engorged females and subsequent individualisation of gravid females for oviposition on a damp filter-paper substrate. Egg batch per female was counted manually using a dissection microscope.

Levels of cytoplasmic incompatibility induction and rescue were measured by crossing various combinations of males and females from the wild-type, *w*AlbA-only and *Wolbachia-negative* lines in pools of 10 males to 5 females, followed by flood-feeding and individualisation as described above. Female mating status post-oviposition was assessed by dissection of spermathecae and visual confirmation of sperm transfer using an inverted 20x compound microscope. The eggs of females lacking visible sperm in spermathecae were discarded from further analysis. Eggs on filter papers from individual females were counted and dried for 5-days, prior to subsequent hatching in water containing bovine liver powder. L1 larval numbers were counted on emergence.

### *Wolbachia* purification and whole-genome amplification

For Illumina sequencing, *w*AlbA was purified by dissecting and pooling ovaries from 27 *w*AlbA-infected *Ae. albopictus* females. Pooled ovaries were rinsed in PBS three times, resuspended in Schneider’s media and homogenized with a sterile pestle. The homogenate was centrifuged at 2,000 g for 2 min to remove cellular debris and the supernatant was filtered through 5 and 2.7 μm sterile filters. The filtrate was centrifuged at 18,500 g for 15 min. The bacterial pellet was resuspended in Schneider’s media and treated with DNase I at 37°C for 30 min to remove host DNA. Following digestion, the DNase was inactivated at 75°C for 10 min and the sample centrifuged at 18,500 g to discard the supernatant. DNA was amplified directly from the bacterial pellet using the REPLI-G Midi kit (Qiagen). The whole-genome amplified DNA (WGA) was cleaned using the QIAamp DNA mini kit (Qiagen). For Oxford Nanopore Technology (ONT) sequencing, ovaries from 100 females were dissected and pooled for DNA extraction as above except that no *Wolbachia* purification or whole-genome amplification was conducted (non-WGA). Finally, DNA was extracted from another pool of 23 pairs of ovaries without WGA to conduct an additional run of Illumina sequencing. However, this sample did not allow us to complete the *w*AlbA genome. Therefore, the corresponding Illumina data was only used for analysing variation in sequencing depth (see below) since reads mapped onto the *w*AlbA genome and plasmids were more evenly distributed than for the WGA sample due to amplification bias.

### Genome sequencing and de novo assembly

A DNA library was prepared from the WGA sample using the Kapa LTP Library Preparation Kit (KAPA Biosystems, Roche7961880001) and sequenced on the Illumina MiSeq platform with the MiSeq Reagent Kit v3 to generate 2×150 bp paired-end reads. The non-WGA sample was used to prepare an ONT library by shearing the DNA into ~8 kb fragments followed by purification and size-selection using AMPure XP beads (Beckman Coulter). The ONT library was then prepared with the Ligation Sequencing Kit (SQK-LSK109) and sequenced for 72 hours using a GridION (ONT) controlled by the MinKNOW software v20.06.9. Illumina and ONT adapters were removed with Trimmomatic v0.38.0 (Bolger et al. 2014) and Porechop v0.2.4 (Wick 2017) respectively. Mosquito reads were filtered out by mapping the Illumina and ONT reads against the *Ae. albopictus* reference assembly (Genbank accession: GCF_006496715.1) using Bowtie2 v2.4.2 (Langmead and Salzberg 2012) and Minimap2 v2.23 (Li 2018) respectively. Unmapped reads were then assembled using the Unicycler hybrid assembly pipeline (Wick et al. 2017). Circular genome maps were created in DNAPlotter (Carver et al. 2009).

### Endpoint PCR and quantitative PCR

Genomic DNA was extracted by homogenizing individual female mosquitoes in 200 μL STE buffer with a sterile pestle, followed by a 30 min incubation at 65°C with 2 μL of Proteinase K (20 mg/mL) and a final incubation for 10 min at 95°C. The extracted DNA was diluted 1/5 in water and samples were centrifuged at 2,000 g for 2 min before PCR. For each PCR reaction, 2 μL of DNA template was amplified using the 2x Taq master mix (Vazyme) in a 25 μL reaction: 12.5 μL of master mix, 12.5 μL of water, 1 μL of each 10 μM primer (Table S5) and the following PCR cycles: 95°C for 3 min, 35 cycles of 15s denaturation at 95°C, 15s for primer annealing (see temperatures in Table S5), 1 min extension at 72°C and a 5 min final extension step at 72°C. *Wolbachia* density relative to host DNA and plasmid copy number were measured by qPCR using the QuantiNova SYBR Green PCR kit (Qiagen) in 10 μL reactions: 5 μL of master mix, 2 μL of water, 0.5 μL of each 5 μM primer (Table S5) and the following cycles: 95°C for 15 min, 40× cycles of 95°C for 15 s and 60°C for 20 s, followed by a melt-curve analysis.

### Search of plasmid sequences in publicly available sequencing data

The presence of plasmid sequences in publicly available *Wolbachia* genomes was searched with TBLASTN (Altschul et al. 1990) using amino acid sequences of all pWALBA1 and pWALBA2 genes. Default parameters and an e-value threshold of 0.05 were used. TBLASTN hits were then visually inspected in Artemis (Carver et al. 2012). When raw sequencing data was available from the Sequence Read Archive (https://www.ncbi.nlm.nih.gov/sra), we attempted to circularize plasmid-like sequences by conducting de novo assemblies using Unicycler. In the case of the *w*Con plasmid, circularization was achieved by visualizing the final assembly graph in Bandage (Wick et al. 2015) and removing an ambiguous path that linked the plasmid to a short multicopy transposase sequence also present in the *Wolbachia* genome (40x sequencing depth relative to the bacterial chromosome). Circular plasmid sequences were then manually rotated to start from the *ParA*-like plasmid partition gene. pWALBA1 and pWALBA2 amino acid sequences were also individually used as queries in the BLASTP online tool (https://blast.ncbi.nlm.nih.gov/Blast.cgi; last accessed January, 2022) to find additional plasmid-like sequences in genome assemblies not annotated as *Wolbachia*. Homologues of the *cif* genes in *Wolbachia* and other *Rickettsiales* genomes were also detected using TBLASTN and representative sequences of the five *cif* types as explained in (Martinez et al. 2021).

In order to investigate gene function and synteny conservation between plasmid sequences, all sequences were reannotated using the RAST annotation pipeline (Aziz et al. 2008) and putative pseudogenes were manually corrected in Artemis. The function of the predicted coding sequences was also inferred from BLASTP searches and using the HHpred webserver (Söding et al. 2005) against the following databases: SCOPe70 (v2.07), Pfam (v35), SMART (v6.0), and COG/KOG (v1.0). Annotated plasmid sequences were then plotted side-by-side along with BLASTN similarities using the R package GenoPlotR (Guy et al. 2010).

Finally, plasmid copy number relative to the *Wolbachia* genome was estimated by mapping the raw sequencing reads onto their respective genome assembly using Bowtie2 v2.4.2 (Langmead and Salzberg 2012) and extracting the sequencing depth values at each position with the Samtools depth command (Li et al. 2009). Sequencing depth per 500 bp windows was calculated and visualized in the R software with a custom script (R Core Team 2013).

### Phylogenetic analysis

The *Wolbachia* phylogeny was generated with RAxML v7.7.6 (Stamatakis 2014) based on the concatenated nucleotide alignment of 36 core genes selected from a set of orthologous single-copy genes described in (Comandatore et al. 2013). For the plasmid and *cif* genes, nucleotide sequences were aligned with MAFFT using the codon-aware tool of the TranslatorX webserver (Abascal et al. 2010) after manually removing premature stop codons from pseudogene sequences. Poorly aligned regions were then trimmed from the alignments using trimAl v1.3 with the –automated1 method (Capella-Gutiérrez et al. 2009). PhyML v3.0 (Guindon et al. 2010) was used to build gene phylogenies with the GTR GAMMA substitution model and 100 bootstrap replicates. Trees were annotated in the iTOL online tool (Letunic and Bork 2007) and the cophylo function from the phytools R package (Revell 2012) for the plasmid and *cif* genes respectively.

## Acknowledgements

This work was supported by the Wellcome Trust (grant numbers 202888, 108508). The raw sequencing reads (accession TBA) and the assembled *w*AlbA genome and plasmids (accessions: CP099802, CP099803, CP099804) are available from the NCBI Genbank database under Bioproject PRJNA852408.

## Supplementary material

**Figure S1. *w*AlbA tissue tropism and fitness effects.** (A) *w*AlbA relative density within tissues of individual females. (B) Number of eggs laid by individual females. (C) Egg hatch rates in the progeny of individual females. Blue, red and grey indicate doubly-infected, *w*AlbA-infected and *Wolbachia*-free mosquito lines respectively. NS: non-significant; *: p < 0.05; **: p < 0.01; ***: p < 0.001.

**Figure S2. Endpoint PCR targeting *w*AlbA and its associated plasmids.** DNA was extracted from individual female mosquitoes and from the Aa23 cell line (n = 4 per *Wolbachia* infection status). Nomenclature: host background-*Wolbachia* infection status. NC: PCR negative control. L: DNA ladder.

**Figure S3. Quantitative PCR analysis of *w*AlbA plasmid copy numbers.** DNA was extracted from individual *w*AlbA-infected female mosquitoes at different timepoints. Females were blood-fed at 8 and 18 day-old. (A) Copy number of a plasmid-specific target (*DnaB*-like gene and Fic family protein for pWALBA1 and pWALBA2 respectively) relative to *Wolbachia* 16S rRNA copies. The *p*-values were calculated with a *t* test on paired log-transformed data. (B) Correlation between *w*AlbA densities and plasmid copy number per mosquito (blue: pWALBA1, red: pWALBA2). The dashed lines show predicted values from linear regressions and *r* is the Pearson’s correlation coefficient.

**Figure S4. Putative chimeric spider contigs**. *A. araneus* (A) and *T. clavipes* (B) contigs were blasted against their respective plasmid-like region, the *w*Mel (Supergroup A) and *w*AlbB (Supergroup B) reference genomes. BLASTN hits were visualized in Bandage. Sequencing depth was measured by mapping the Illumina and Nanopore reads from the corresponding sample onto the contig of interest.

**Figure S5. Phylogenies of *ParA*-like, *ParG*-like, HP1 and *DnaB*-like homologues.** Full circles indicate the genomic location of the different homologues. Branch support calculated from 100 bootstraps replicates and >80% are shown.

**Figure S6. Phylogenies of phage portal, terminase large subunit, stabilization and coat protein homologues.** Full circles indicate the genomic location of the different homologues. Branch support calculated from 100 bootstraps replicates and >80% are shown.

**Figure S7. Phylogenies of HP2, HP3, HP4 and *QueT*-like transporter protein homologues.** Full circles indicate the genomic location of the different homologues. Branch support calculated from 100 bootstraps replicates and >80% are shown.

**Figure S8. Phylogenies of HP7, HP9, PD-(D/E)XK nuclease and Fic/DOC protein homologues.** Full circles indicate the genomic location of the different homologues. Branch support calculated from 100 bootstraps replicates and >80% are shown.

**Figure S9. Phylogenies of *RelB* and *RelE* homologues.** Full circles indicate the genomic location of the different homologues. Branch support calculated from 100 bootstraps replicates and >80% are shown.

**Figure S10. Association of *cif* genes with Integrative Conjugative elements.** Similarity is indicated by gene colours (BLASTP) and by the grey areas between sequences (TBLASTX) where darker grey means more similar. Dark grey genes are transposable element sequences.

**Table S1. *Wolbachia* genome statistics.**

**Table S2. *w*AlbA prophage regions.**

**Table S3. Plasmid gene BLASTP hits and HHpred predicted domains.**

**Table S4. List of *Wolbachia* genome assemblies and detected plasmid-like regions.**

**Table S5. List of PCR and qPCR primers.**

