## Supplementary figures and images for "Genome sequencing and comparative analysis of *Wolbachia* strain *w*AlbA reveals *Wolbachia*-associated plasmids are common"

### Figure S1

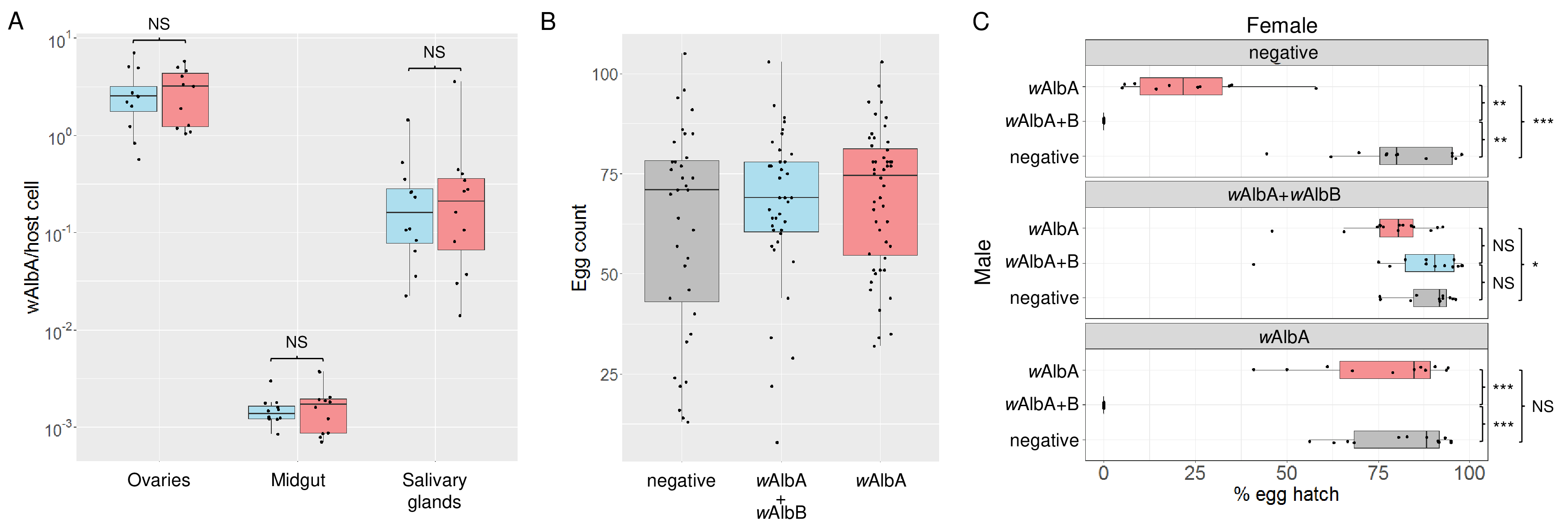

### Figure S2

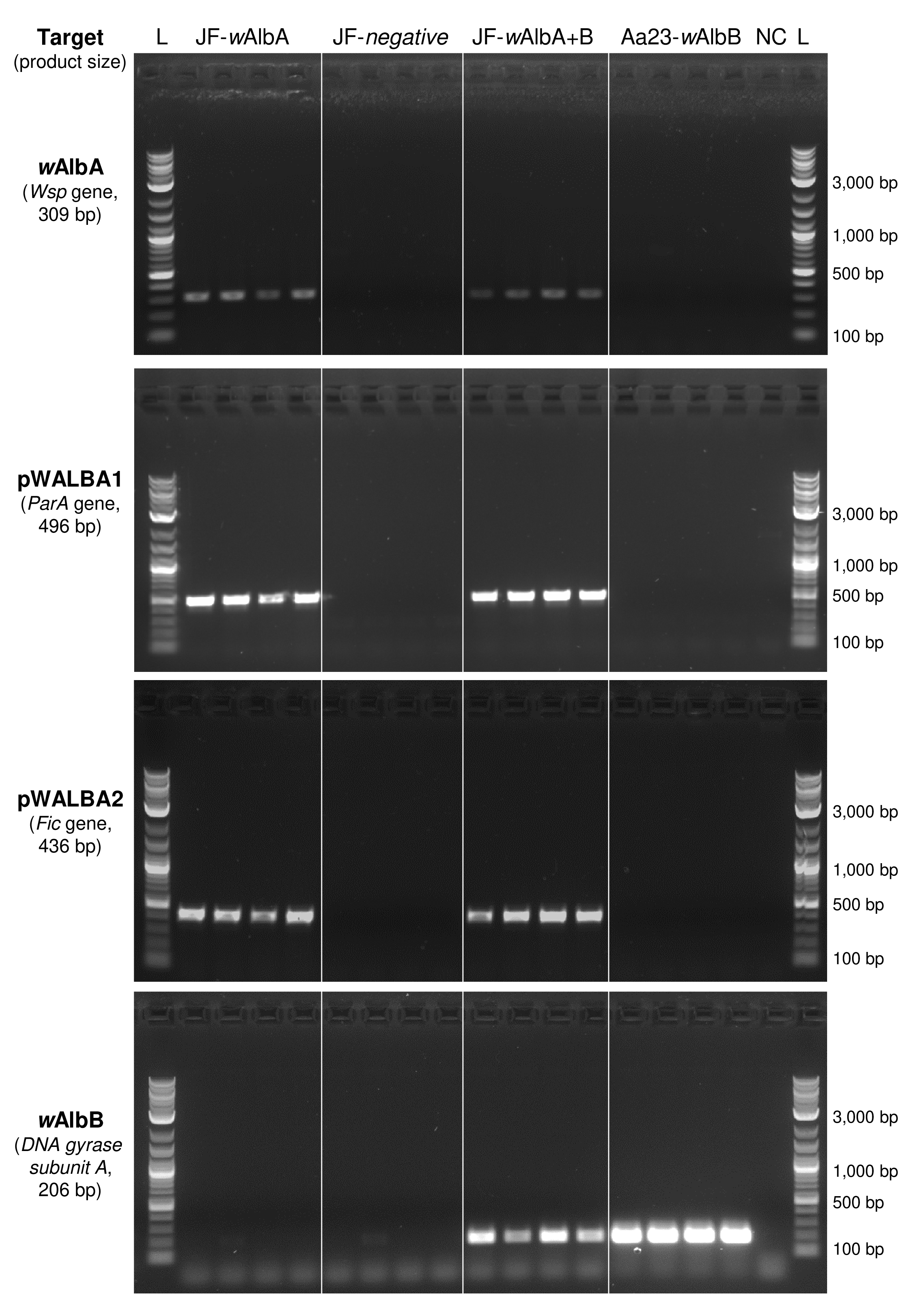

### Figure S3

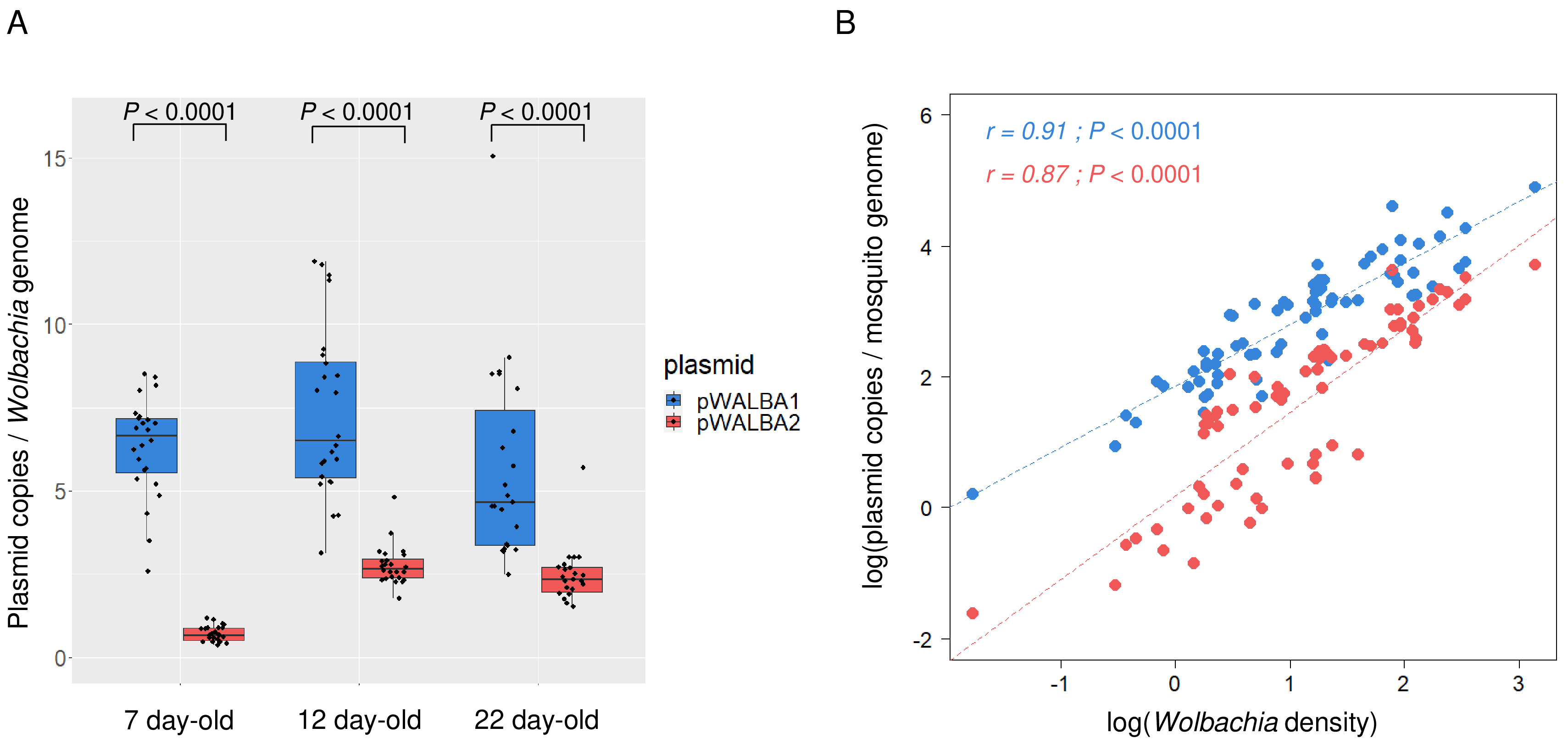

### Figure S4

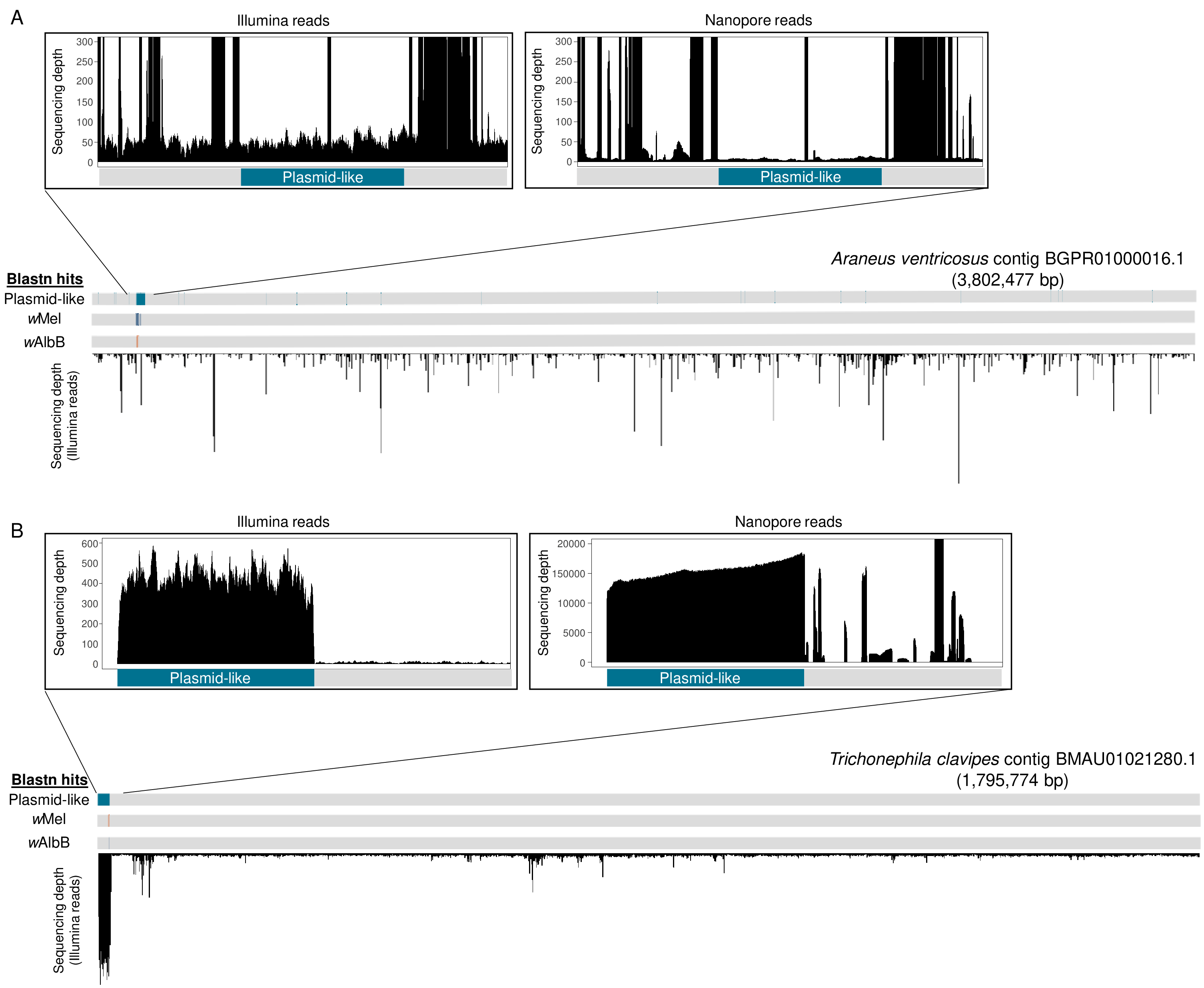

### Figure S5

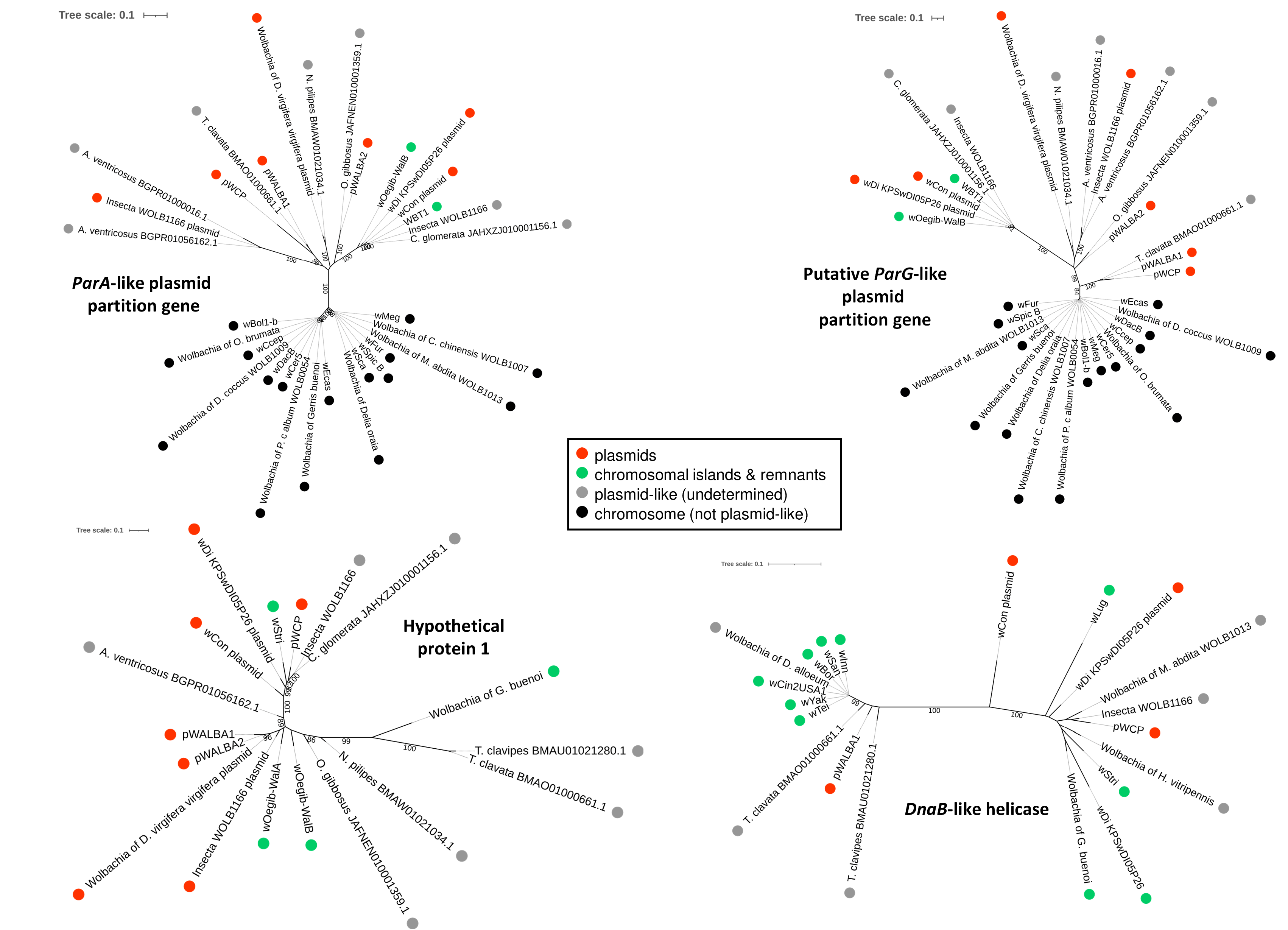

### Figure S6

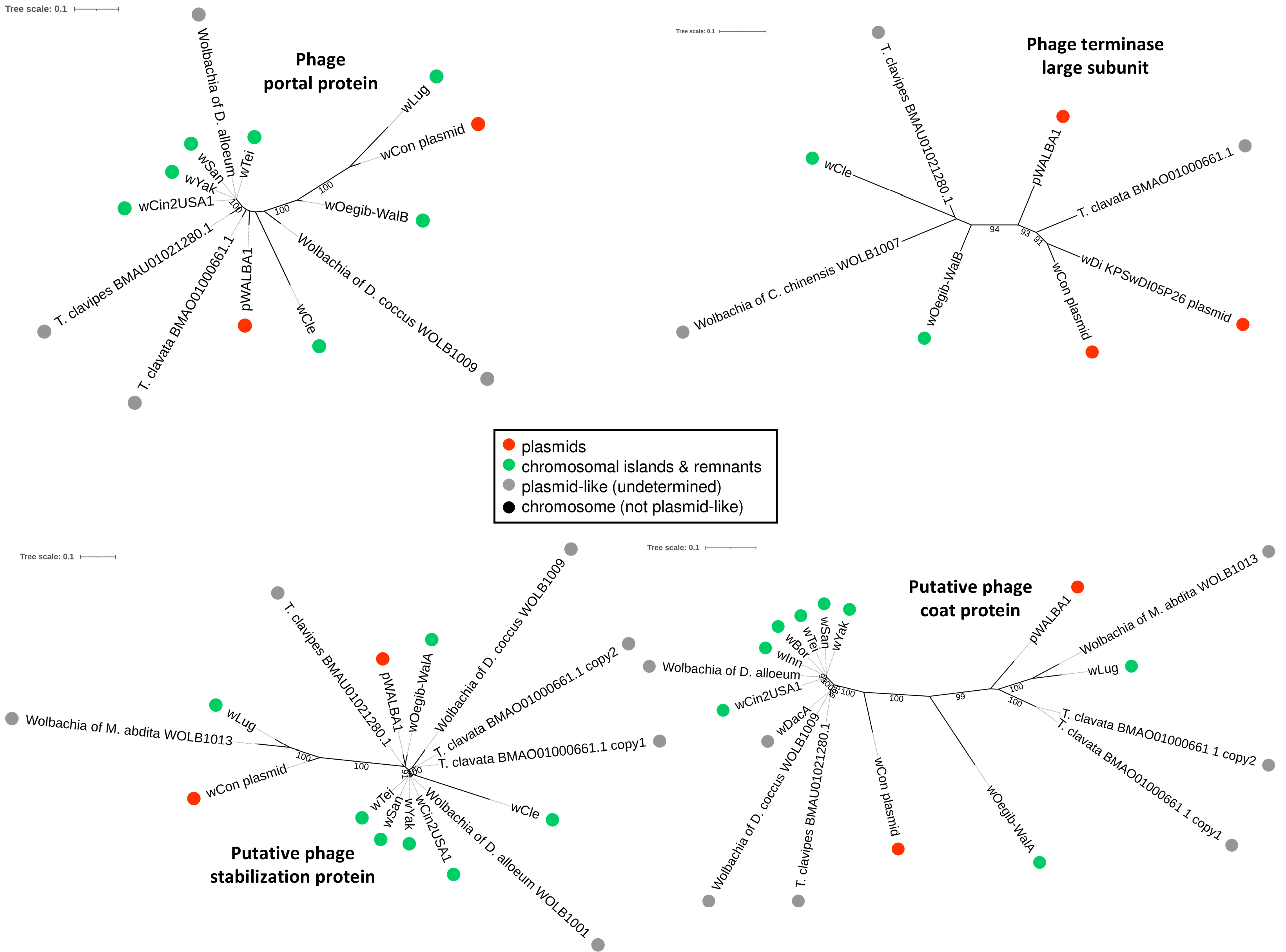

### Figure S7

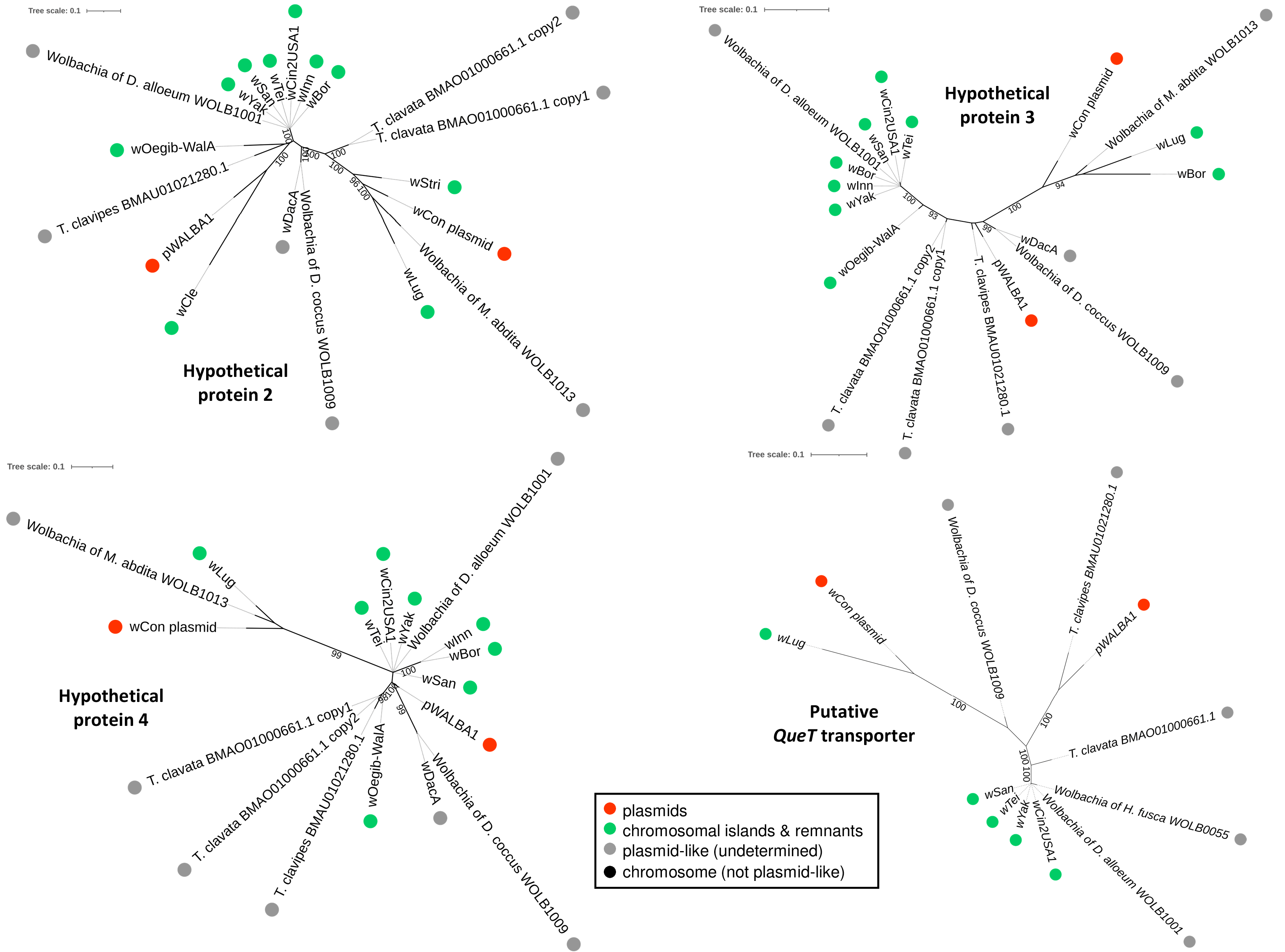

### Figure S8

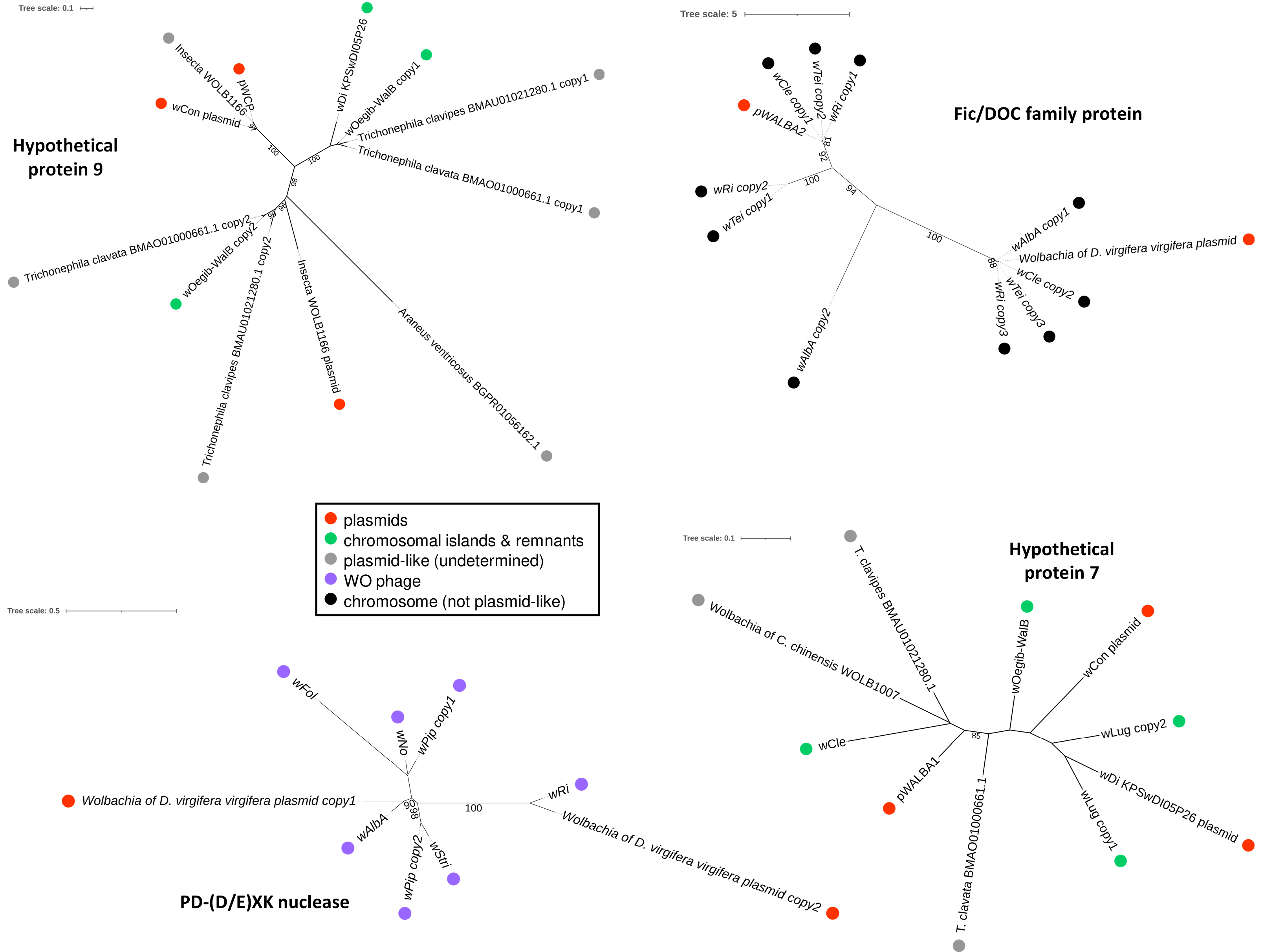

### Figure S9

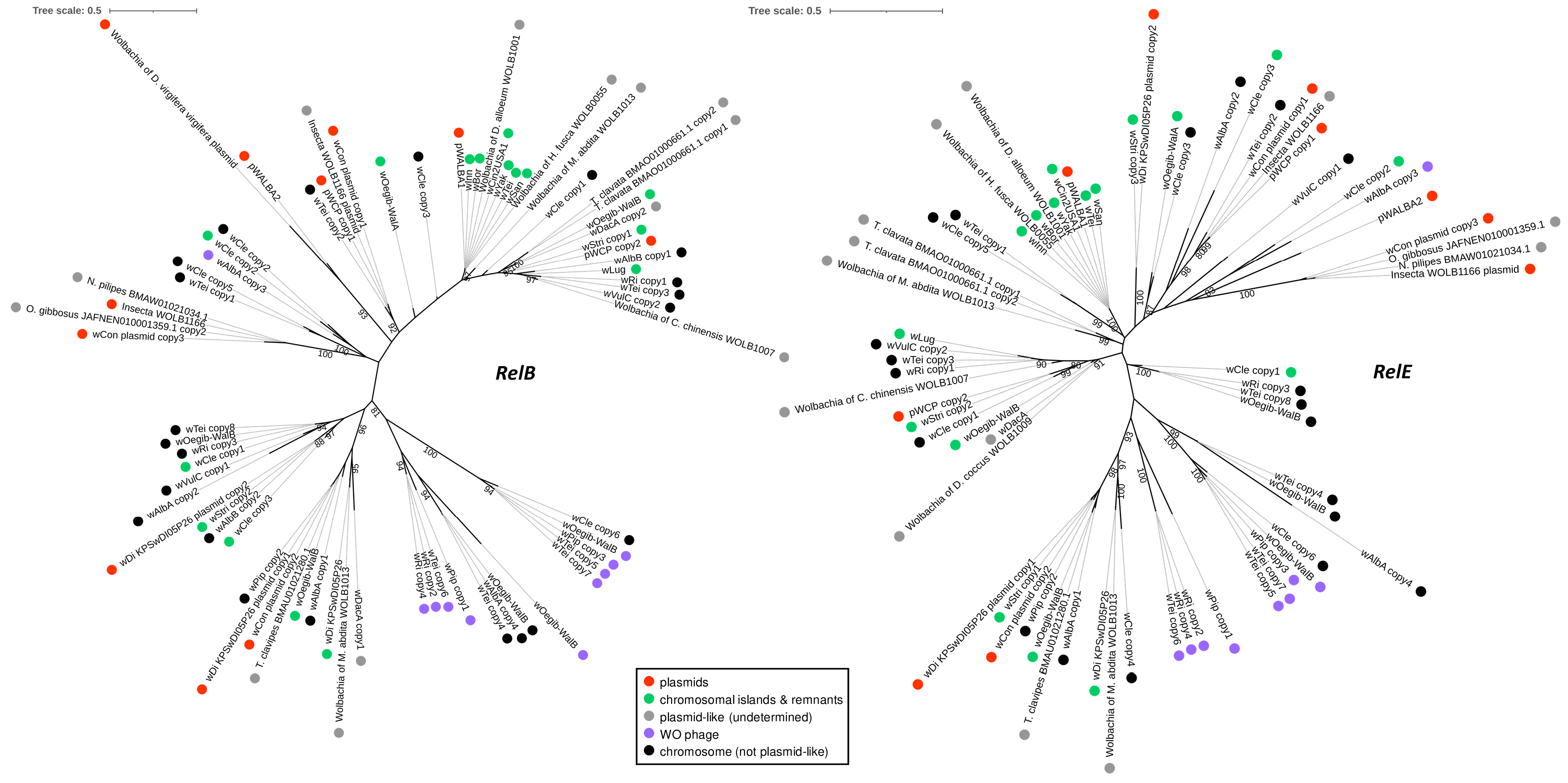

### Figure S10

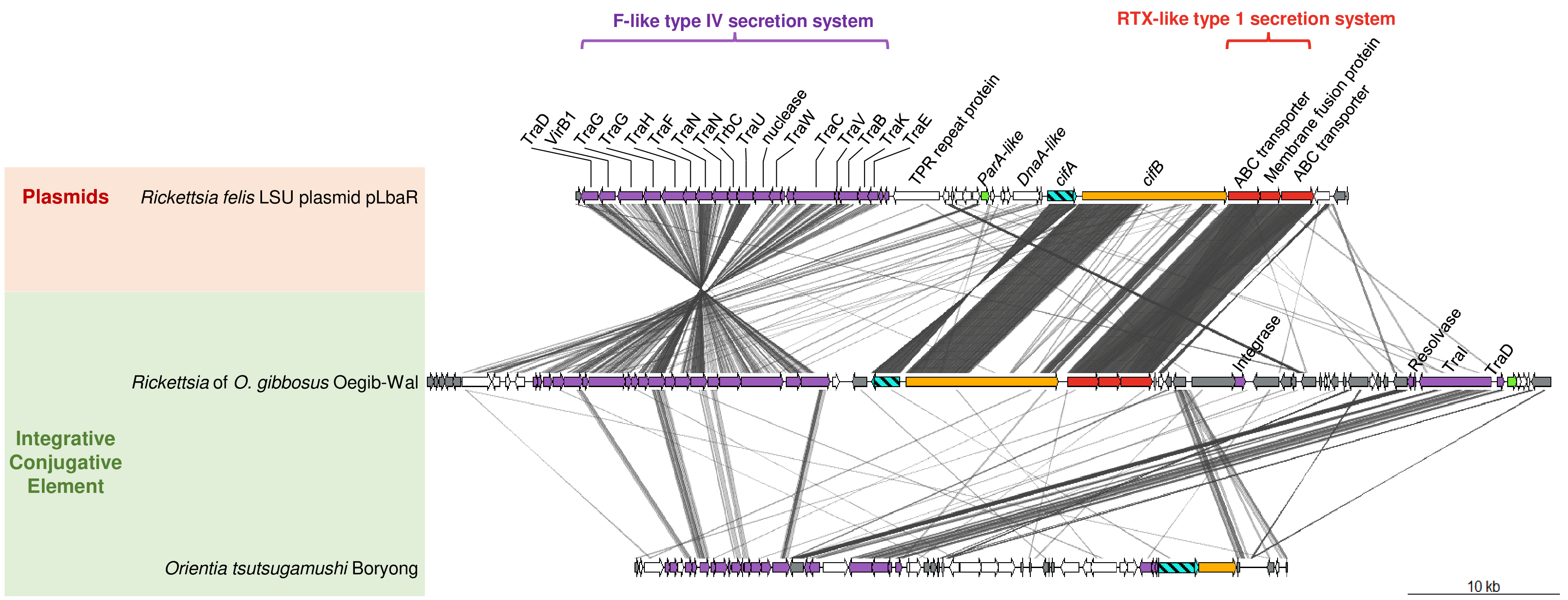
